# A new efficacious Mcl-1 inhibitor maximizes bortezomib and venetoclax responsiveness in resistant multiple myeloma cells

**DOI:** 10.1101/2023.12.06.570435

**Authors:** Omar S. Al-Odat, Krishne Gowda, Sandeep K. Srivastava, Shantu G Amin, Subash C. Jonnalagadda, Tulin Budak-Alpdogan, Manoj K. Pandey

## Abstract

Despite a record number of clinical studies investigating various anti-cancer drugs, the 5-year survival rate for multiple myeloma (MM) patients in the United States is only 55%, and nearly all patients relapse. Poor patient outcomes demonstrate that myeloma cells are “born to survive,” which means they can adapt and evolve following treatment. As a result, new therapeutic approaches to combat this survival mechanism and target treatment-resistant malignant cells are required. Mcl-1, an anti-apoptotic protein, is required for the development of MM and resistance to therapy. This study looks at the possibility of KS18, a Mcl-1 inhibitor derived from pyoluteorin, to treat resistant MM. We show that KS18 inhibits Mcl-1 selectively and promotes post-translational modifications, resulting in UPS-dependent Mcl-1 degradation. Our findings show that KS18-induced Mcl-1 degradation results in caspase-dependent apoptosis. Importantly, KS18 triggered apoptosis in MM patient samples and bortezomib-resistant cells, synergizing with venetoclax to boost apoptosis. Furthermore, KS18 inhibits colony formation in bortezomib-resistant cells. KS18 treated NSG mice displayed significant tumor shrinkage without significant toxicity after four weeks of therapy with a single acceptable dose each week, indicating its powerful anti-neoplastic and anti-resistance characteristics. This study strongly implies that KS18 may treat MM and provide new hope to patients who are experiencing recurrence or resistance.

**Key points:**

- Given that KS18 is a robust Mcl-1 inhibitor that targets Mcl-1 efficiently, it has the potential to be a novel treatment for multiple myeloma.
- KS18 has shown promise in re-sensitizing myeloma cells to chemotherapy as well as in overcoming resistance to bortezomib, venetoclax, and ABT-737.

## INTRODUCTION

Multiple myeloma (MM) is a bone marrow-based hematologic cancer caused by clonal plasma cell growth.^1,2^ These cells have many cytogenetic abnormalities that affect prognosis and treatment.^3^ A proteasome inhibitor, dexamethasone, and immunomodulatory or chemotherapeutic drug regimen with or without autologous stem cell transplantation is common treatment.^4–6^ These drugs are effective and often used to support other treatments, but intrinsic or acquired drug resistance makes myeloma incurable, and all patients recur.^7^

The expression of anti-apoptotic members of the Bcl-2 family is a critical factor in myeloma cell survival.^8^ The presence of anti-apoptotic (e.g., Bcl-2, Bcl-xL, Mcl-1) and pro-apoptotic (e.g., Bax, Bak, Bim, Puma, Bid, Noxa) proteins determines whether myeloma cells undergo apoptosis. Mcl-1 expression, in particular, is required for MM cell survival.^9–11^ Indeed, anti-sense RNA suppression of Mcl-1 causes death in myeloma cells, whereas targeting Bcl-2 or Bcl-xL had little or no effect.^9^ Furthermore, a considerable threshold level of Mcl-1 is required for myeloma cell viability *in vitro*, whereas clinically, overexpression of Mcl-1 is detected in 52% of MM patients at diagnosis and 81% at relapse, implying that Mcl-1 level corresponds with disease progression.^12^ The same clinical trials discovered that increased Mcl-1 expression is related with a shorter lifespan.^12^

In addition to Mcl-1, MM includes overexpression of Bcl-2 and/or Bcl-xL, suggesting these 3 prosurvival proteins are attractive therapeutic targets.^13,14^ However, Mcl-1, Bcl-xL, and Bcl-2 expression alone cannot explain Bcl-2 family dependency. Instead, Bim’s preference for Mcl-1 over Bcl-2 or Bcl-xL dictates the medicines’ fate in Mcl-1-expressing myeloma cells.^15^ Drugs that mimic the Bcl-2 homology 3 (BH3) domains of pro-apoptotic Bcl-2 family members to neutralize these proteins by attaching to their surface hydrophobic grooves are an interesting therapy option. ABT-199 (venetoclax) is the first FDA-approved Bcl-2-specific BH3 mimetic for 17p chromosomal deletion chronic lymphocytic leukemia patients.^16^ MM patients with a (11;14) translocation [t(11;14)] express more Bcl-2 than Bcl-xL or Mcl-1 and respond well to venetoclax alone.^17,18^ MM cells with high Mcl-1 expression are less sensitive to venetoclax,^13^ but downregulation can overcome this. ^19–24^ These findings show that earlier therapies or clonal selection may have shifted cellular plasticity toward Mcl-1 dependency during disease development.^25^

The research shows that Mcl-1 contributes to treatment resistance and illness progression. Mcl-1-targeted therapy may benefit MM patients, especially those in relapse. Unfortunately, no FDA-approved drug directly targets Mcl-1. Several selective Mcl-1 inhibitors are being developed or evaluated in human trials, although earlier trials were halted due to cardiac toxicity (NCT03218683), (NCT03797261), (NCT03465540), and (NCT03218683).^26–28^ To address these limitations, our team created KS18, a pyoluteorin derivative Mcl-1 inhibitor, to test its recurrence-prevention potential.^29^ KS18 is an effective Mcl-1 inhibitor, enhances other chemotherapeutic drugs, and re-sensitizes chemotherapy-resistant MM cells. We proposed using KS18 to find new MM therapies, especially chemotherapy-resistant ones.

## MATERIALS AND METHODS

### Patient samples, cell lines, and reagents

Primary human MM samples from newly diagnosed patients were obtained from Penn State University Institute biobank. Bortezomib-resistant cells were generously donated by Dr. Nathan Dolloff, Medical University of South Carolina, Charleston, SC.^30^ By dose escalation, we developed resistant MM cells against venetoclax, ABT-737, and lenalidomide (U266-VEN-R, U266-ABT-R, and U266-LEN-R).^31^ MM.1S, MM.1R, U266, and RPMI 8226 human MM cells were procured from ATCC (Manassas, VA) and grown in respective medium using our laboratory’s protocol.^32^ KS18 agent was synthesized and characterized at Penn State College of Medicine, Hershey, PA.^29^ All other agents were procured from Selleckchem (Houston, TX).

### Cell viability assay

MTT assay measured cell viability. The MTT assay was carried out essentially as described previously.^32^

### Western blot

Western blotting was performed essentially as described before.^32^ Using Gentle ReView™ Stripping Buffer (VWR, #19G0856497), several protein detections were done on the same membrane to conserve time and samples. We repeated all crucial blot experiments two to three times.

### Immunoprecipitation (IP)

Immunoprecipitation followed by western blotting was performed with either MCl-1 or Bim antibodies by following our laboratory protocol as described before.^19^

### Cytochrome C release test

The human MM cells were treated with KS18 and incubated for 24h at 37^0^C. Following incubation, the cytosolic fraction was extracted using the cytochrome C release assay kit (Abcam, #ab65311) according to the manufacturer’s instructions. Protein quantification and western blotting were carried out as previously described.^32^

### Annexin V live dead assay

Human MM cells were plated, treated, and incubated for 24h, and then tested for Annexin V Live Dead assay (Luminex Corporation, #MCH100105). Cells were stained with 1:1 Annexin V at room temperature in the dark for 20 minutes following manufacturer’s protocol. Data was analyzed using Muse® Cell Analyzer.^32^

### Caspase-3/7 assay

Human MM cells were treated and incubated for 24 hours before being tested for caspase 3/7 (Luminex Corporation, # MCH100108). Cells were stained for 30 minutes at 37°C with caspase 3/7 working solution following manufacturer’s protocol. The 7-AAD solution was added after incubation. Data was analyzed using Muse® Cell Analyzer.^32^

### Flow cytometry

MM patient samples or human MM cells were plated, treated, and incubated for 24h before being tested using Annexin V Apoptosis Detection Kit (BD Biosciences, #556547). Annexin V/PI stain solution was combined with cells and incubated at room temperature in the dark for 20 minutes before detection of dead cells using flow cytometer (BIO-RAD, Se^TM^ Cell Sorer).^32^

### Immunostaining and confocal microscopy

Immunostaining was performed essentially as described before.^32^ Photographed was done by using a Nikon A1R GaAsP Laser Scanning Confocal Microscope (Nikon Eclipse Ti inverted with a 60x objective).

### Colony forming assay

Colony forming assay was performed essentially following the manufacturer’s protocol. Briefly, human MM cells were cultured at a density of 1 × 10^4^ cells in MethoCult™ H4230 media (Stem Cell Technologies, #04230) supplemented with 15% FBS and 10% Gibco Phytohemagglutinin M form (PHA-M) (Thermo Fisher, #10576015) stimulated leukocyte conditioned medium. Human MM cells were incubated at 37°C in 5% CO2 for 14 days before colonies were visually counted using a microscope.

### Toxicity and efficacy studies in mice xenograft model

Female NOD-SCID-IL2R gamma null (NSG) mice aged five weeks were procured from Jackson Laboratories and maintained and monitored in the animal research facility at Cooper Medical School of Rowan University (CMSRU), Camden, NJ. All xenograft investigations were conducted at CMSRU in accordance with the ethical rules provided by Rowan University’s own Institutional Animal Care and Use Committee (IACUC). Mice were subcutaneously injected into the right flank with 5 X10^6^ U266 MM cells in a 1:1 ratio of RPMI-1640 and Matrigel basement membrane matrix (Becton Dickinson, Bedford, MA) to create a human MM xenograft. When palpable tumors (volume ∼100 mm^3^) appeared about 10 days following injection, animals were randomly assigned into four groups of five mice each. To quantify tumor volume, digital caliper measurements of the longest perpendicular tumor diameter were taken as follows: 4/3 X (width/2)^2^ X (length/2). Every three days, the tumor volume and body weight were measured. The animals were euthanized at the end of the study, and the tumors were removed, weighed, and snap-frozen for future research.

### Statistical analysis

Statistical significance of the results was analyzed by unpaired two-tailed Student’s t-test or one-way analysis of variance (ANOVA), or two-way ANOVA using GraphPad prism software. Each graph represents the average of at least three replicates with error bars on the graph representing standard deviation* P ≤ 0.05, ** P ≤ 0.01, *** P ≤ 0.001, **** P ≤ 0.0001. ns: non-significant.

## RESULTS

### KS18 induces the degradation of Mcl-1 in MM cells

We used X-ray crystal structure of Mcl-1 from the Protein Data Bank (PDB ID: 2JM6, 2.5 Å) to perform docking studies using Autodock Vina program. Docking results suggest that KS18 interacts with Mcl-1.^34^ The protein-protein restraint docking of NOXA with Mcl-1 using the HADDOCK (high ambiguity-driven protein-protein docking) algorithm confirmed that KS18 binds Mcl-1 at its NOXA binding sites (Supplementary Fig. 1A).{Warren, 2019 #57} We next determined the expression of anti-apoptotic family members in MM cell lines. Most myeloma cells, except RPMI 8226, overexpressed Mcl-1, suggesting these cells are dependent on Mcl-1 for survival. In contrast, RPMI 8226 cells are likely to be depend on Bcl-2 and Bcl-xL (Supplementary Fig. 1B). We further investigated the effect of KS18 on Mcl-1. As shown in Fig. 1A&B, KS18 inhibits Mcl-1 expression in a dose- and time-dependent manner, but importantly, did not demonstrate any effect on Bcl-2/Bcl-xL expression. Similarly, KS18 was effective at lowering Mcl-1 expression in two additional MM cell lines (MM.1S, and MM.1R; Supplementary Fig. 1C). Immunocytochemical analysis also suggested that KS18 inhibits Mcl-1 expression in U266 MM cells (Fig. 1C). Mcl-1 degradation is controlled by phosphorylation at the Ser159/Thr163 sites, which leads to ubiquitination by E3 ligases like F-box and WD repeat domain-containing 7 (FBW7), Mule, and b-TrCP.^35^ To understand how KS18 degrades Mcl-1, we tested the effect of KS18 on the phosphorylation and ubiquitination of Mcl-1, as shown in Fig. 1D, KS18 induces the phosphorylation of Mcl-1 at Ser159/Thr163 residues, followed by ubiquitination (Fig. 1E). We found that phosphorylation event was maximum within 6h of the treatment followed by the degradation of Mcl-1 starting from 3h (Supplementary Fig. 1D).

**Figure 1:**
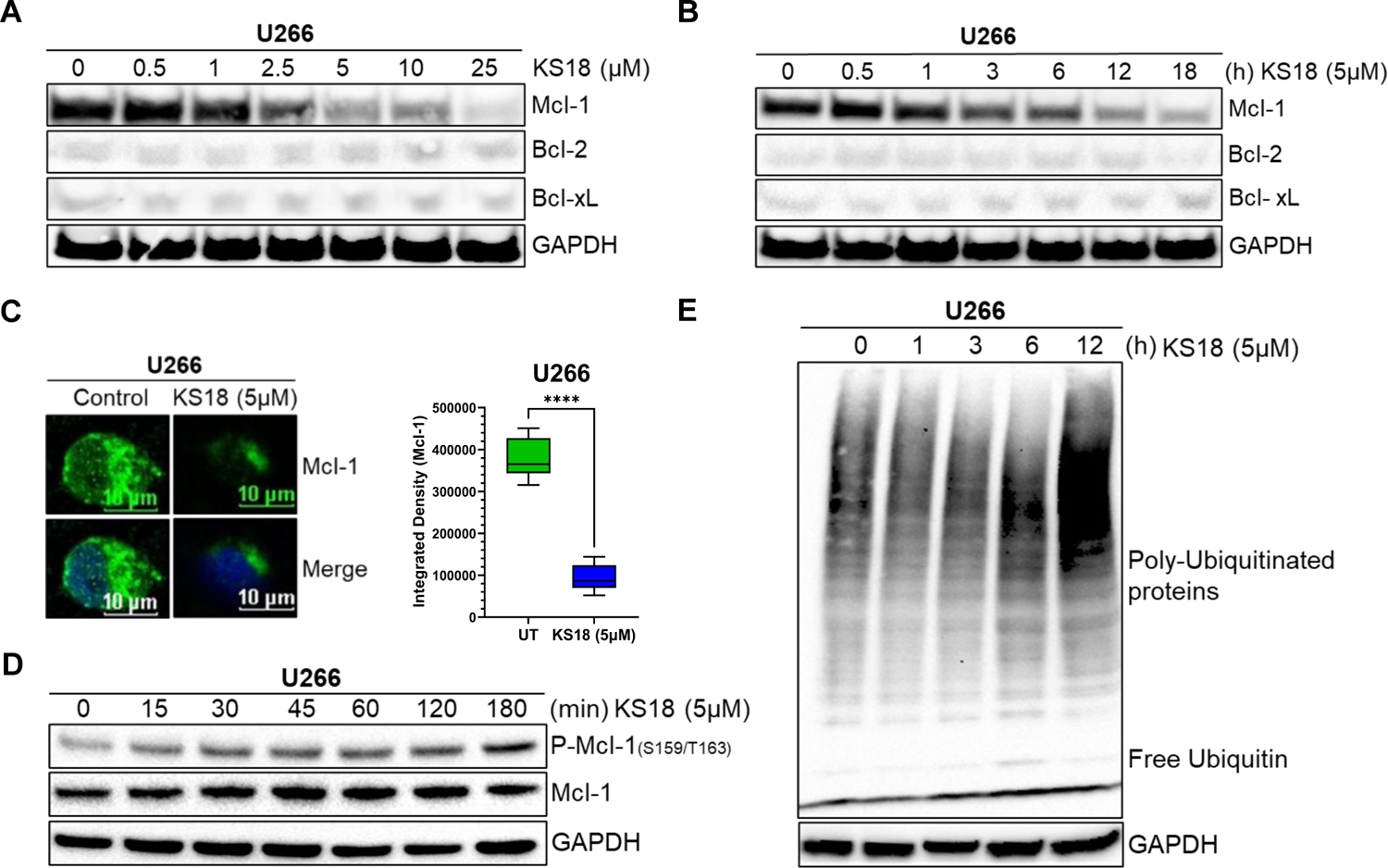
Effect of KS18 on Mcl-1 in MM cells. *A-B*, KS18 suppresses Mcl-1 in a dose- and time-dependent manner. U266 MM cells were treated with the indicated doses of KS18 for 24h or with 5μM at various time points. After incubation, the cells were lyzed and immunoblotted with the aforementioned antibodies. GAPDH antibody was used as a loading control on the stripped membrane. ***C,*** Immunocytochemical study of Mcl-1 using confocal microscopy. For 24h, cells were treated with 5μM of KS18. Following incubation, immunocytochemical examination was carried out as indicated in the ‘MATERIALS AND METHODS’ section. ImageJ was used to calculate the integrated density of cells. **** P ≤ 0.0001. ***D&E,*** Mcl-1 was phosphorylated and ubiquitinated after being exposed to KS18. 5μM of KS18 was applied to U266 MM cells at various time points. After incubation, the cells were lyzed and immunoblotted with the aforementioned antibodies. GAPDH antibody was used as a loading control on the stripped membrane.

Together these results demonstrate that treatment of KS18 induces the phosphorylation of Mcl-1 as early as 15 minutes followed by its degradations, suggesting that KS18 induces ubiquitin/proteasome-dependent protein degradation (UPS) of Mcl-1 (Fig. 1D-E; Supplementary Fig. 1D).

### KS18 induces apoptosis in a caspase-dependent manner

A network of pro- and anti-apoptotic proteins controls cell destiny (Fig. 2A). BH3 domains of pro-apoptotic proteins bind to anti-apoptotic proteins’ hydrophobic groove. In Mcl-1-expressing myeloma cells, Bim distribution dictates Mcl-1 dependency or codependence with Bcl-2/Bcl-xL.^36,37^ Mcl-1’s binding to pro-apoptotic protein Bim was determined. Fig. 2B shows that KS18 lowered Bim_EL_ but not Bim_L_ or Bim_S_. Bid expression was likewise decreased in KS18-treated cells. Using immunoprecipitation, we pulled down Mcl-1 and Bim to explore their relationship. In KS18-treated cells, reduced Mcl-1 expression was related to poor binding of Bim_EL_ followed by Bax (Fig. 2C). We also found similar results when we pulled down Bim followed by Mcl-1 immunoblotting, demonstrating that Bim dissociates from Mcl-1 and activates Bax. Therefore, Bax and Bim are the major mediators of KS18-induced apoptosis via Mcl-1 degradation. We examined KS18’s impact on Noxa protein. KS18 slightly affected Noxa expression (Fig. 2E). Bax and Bak oligomerize and produce pores in the outer mitochondrial membrane, releasing cytochrome C and other apoptogenic factors.^38^ KS18 therapy activates Bax (Fig. 2E), thus we explored if it releases cytochrome C. KS18 treatment released cytochrome C into the cytosol (Fig. 2F). In U266 cells, KS18 stimulates caspase-3 cleavage via procaspase-3 processing and PARP cleavage (Fig. 2G). We verified this in two additional MM cell lines, MM.1R and MM.1S. KS18 processes procaspase-3 and cleaves PARP in both cells (Supplementary Fig. 1E). KS18’s apoptosis-inducing capacity in MM cells was examined by caspase-3/7 and Annexin V staining. KS18 substantially causes caspase-dependent MM cell death (Fig. 2H-I; Supplementary Fig. 1F-G). These data suggest that KS18 may induce intrinsic apoptosis in MM cells via promoting Bax-induced mitochondrial permeabilization, cytochrome C release, and caspase activation followed by PARP cleavage.

**Figure 2:**
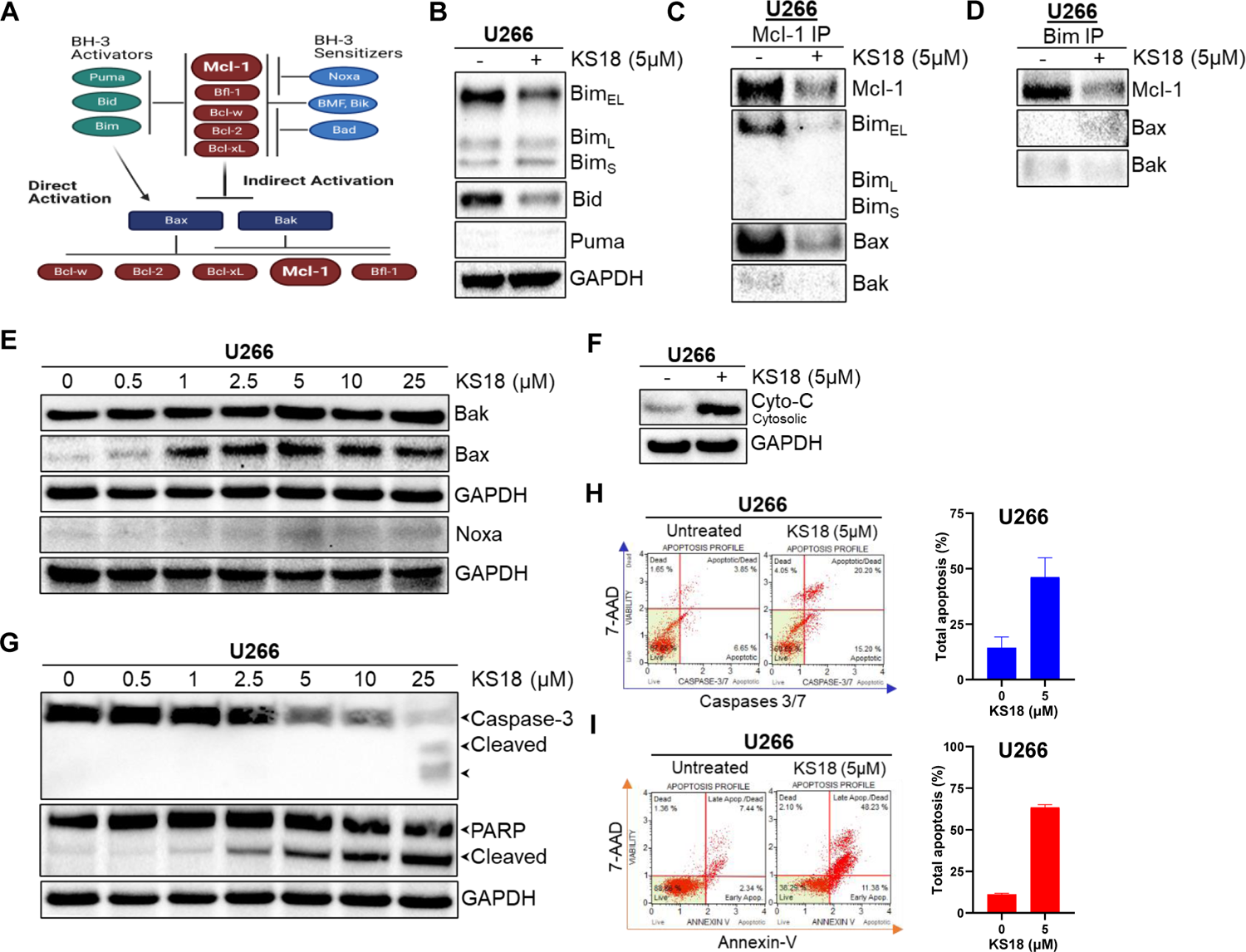
Effect of KS18 on apoptosis regulators. ***A,*** Intrinsic pathway is promoted by cellular stresses that modulate Bcl-2 family proteins and activates Bak and Bax. In the indirect activation, upregulation of BH3-only proteins will act as inhibitors of anti-apoptotic proteins by competing for their binding with Bax and Bak proteins, leading Bax and Bak to oligomerize. In the direct activation, upregulation of BH3 activators proteins directly activates Bax and Bak. ***B,*** the effect of KS18 on BH3 activators proteins (Bim, Bid, and Puma). ***C&D,*** cells were treated with KS18 at the indicated concentration for 24h before being lyzed. As specified in ‘MATERIALS AND METHODS’ section, the whole cell extract was immunoprecipitated with either Mcl-1 or Bim, and immunoblotting was done with the aforementioned antibodies. ***E,*** Dose-dependent effect of KS18 on various apoptosis inducers. ***F,*** effect of KS18 on cytochrome C release in cytoplasm. Cells were treated first with KS18 (5μM) for 24h. The cytosolic fractions were isolated using a cytochrome C kit as described under ‘MATERIALS AND METHODS’ section and subjected to immunoblotting for protein analysis with cytochrome C antibody and anti-GAPDH to assess loading. ***G,*** Dose-dependent effect of KS18 on caspase-3 and PARP proteins in U266 cells. For sections B, E, and G, cells were treated for 24 h with the indicated quantity of KS18. Following incubation, cells were collected, lyzed, and immunoblotting for the aforementioned antibodies was done. The anti-GAPDH blotting was performed to determine loading. ***H&I,*** Caspases 3/7 and Annexin V staining were used to identify apoptotic cells. U266 cells were treated with KS18 (5µM) for 24h before being stained with caspases 3/7 dye or Annexin V dye as described in ‘MATERIALS AND METHODS’ section and evaluated using the Muse® Cell Analyzer. The total number of apoptotic cells was enumerated, and a graph was created using the GraphPad prism software.

### KS18 is highly efficacious as monotherapy and enhances the cytotoxic effects of bortezomib in MM cells

KS18’s cytotoxic capability was examined using the MTT assay and compared to other chemotherapy drugs. KS18 can kill various MM cells over time, as illustrated in Fig. 3A. KS18 outperformed other MM chemotherapeutic drugs (Fig. 3B; Supplementary Fig. 2A-C). Further, we tested its apoptotic efficacy in MM samples. KS18 therapy (5µM for 24h) was effective in MM patient samples, resulting in 88.8% apoptosis (Fig. 3C). We investigated whether KS18 can be used in combination with bortezomib which is a first line therapy. Fig. 3D demonstrates that KS18 enhances the therapeutic efficacy of bortezomib, KS18 (5µM) and bortezomib (5nM) combination reducing MM cell viability by 56-70% (Fig. 3D; Supplementary Fig. 2D).

**Figure 3:**
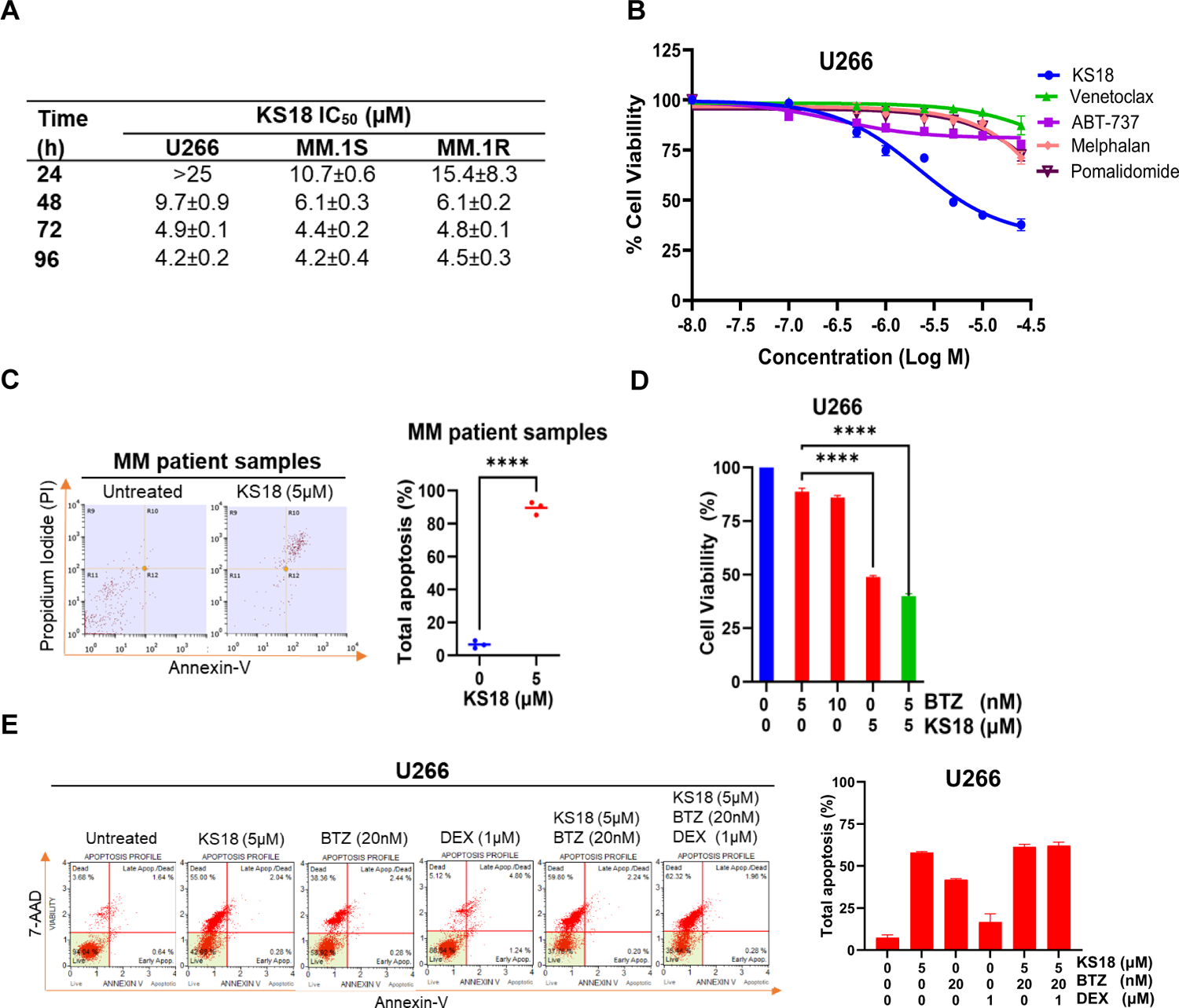
KS18 is effective against MM cells, synergize with bortezomib and causes apoptosis in MM patients. ***A,*** A panel of human MM cell lines (U266, MM.1S, and MM.1R) were treated with increasing doses of KS18 (0-25μM) for 24, 48, 72, and 96h, and the cytotoxicity of KS18 was determined using the MTT test. The IC_50_ values were calculated using GraphPad prism. ***B,*** Cell viability was evaluated using the MTT test after 72 h of treatment with increasing dosages (0-25μM) of KS18 and other chemotherapeutic drugs (venetoclax, ABT-737, melphalan, and pomalidomide). GraphPad prism was used to create the graph. ***C,*** Annexin V/propidium iodide (PI) staining was used to examine KS18’s ability to induce apoptosis in MM patient samples. All MM patient samples were treated for 24h with KS18 (5µM), and the Annexin V Apoptosis Detection Assay was performed using flow cytometry. Total apoptosis was estimated as ([% cell death in treated cells − % cell death in control] / (% viable cells control) ×100, and the graph was created using GraphPad prism. **** P ≤ 0.0001. ***D***, U266 cells were treated for 72h with bortezomib (BTZ) (5 and 10nM) alone and BTZ (5nM) in conjunction with KS18 (5µM), and cell viability was determined using the MTT test. GraphPad prism was used for graphical representations and statistical analysis. **** P ≤ 0.0001. ***E***, Apoptotic cells are detected by Annexin V staining. U266 cells were treated for 24h with KS18 (5µM) alone or in combination with BTZ (20nM) and DEX (1µM), then stained with Annexin V dye and analyzed using the Muse® Cell Analyzer. GraphPad prism was used for graphical representations and statistical analysis.

For standard-risk myeloma patients, the clinic advises dexamethasone and bortezomib or lenalidomide.^39^ Cyclophosphamide, melphalan, or doxorubicin can be added when rescue regimens fail if the sickness is resistant.^40–42^ We evaluated KS18 with dexamethasone and bortezomib for clinical simulation. KS18 in combination with bortezomib and low-dose dexamethasone improves therapeutic response, suggesting that KS18 may improve therapeutic outcomes (Fig. 3E). A similar impact was seen in MM.1S cells (Supplementary Fig. 1E).

### The therapeutic efficacy of Bcl-2/Bcl-xL inhibitors in MM cells can be improved with the addition of the Mcl-1 inhibitor KS18

Although venetoclax is a selective small-molecule Bcl-2 inhibitor, it is ineffective against cancer cells with high Mcl-1 levels, including MM.^43–46^ We tested whether KS18 sensitizes Mcl-1 overexpressing cells to venetoclax by treating MM cells with either KS18 or venetoclax alone or in combination. The treatment of venetoclax alone induces the expression of Mcl-1, but it was suppressed when treated with both KS18 and venetoclax (Fig. 4A). Similarly, this was observed in other MM cells (Supplementary Fig. 3A). Furthermore, the apoptosis-inducing potential of KS18 in combination with venetoclax was investigated. In U266 cells, the addition of KS18 (5µM) to venetoclax (0.5µM) exhibited a strong additive impact, with significant stimulation of caspase-mediated apoptosis (Fig. 4B-C). The same response was observed in MM.1S cells (Supplementary Fig. 3B-C). Furthermore, the addition of KS18 (5µM) to venetoclax resulted in a considerable reduction in cell viability in MM cells (Supplementary Fig. 3D).

**Figure 4:**
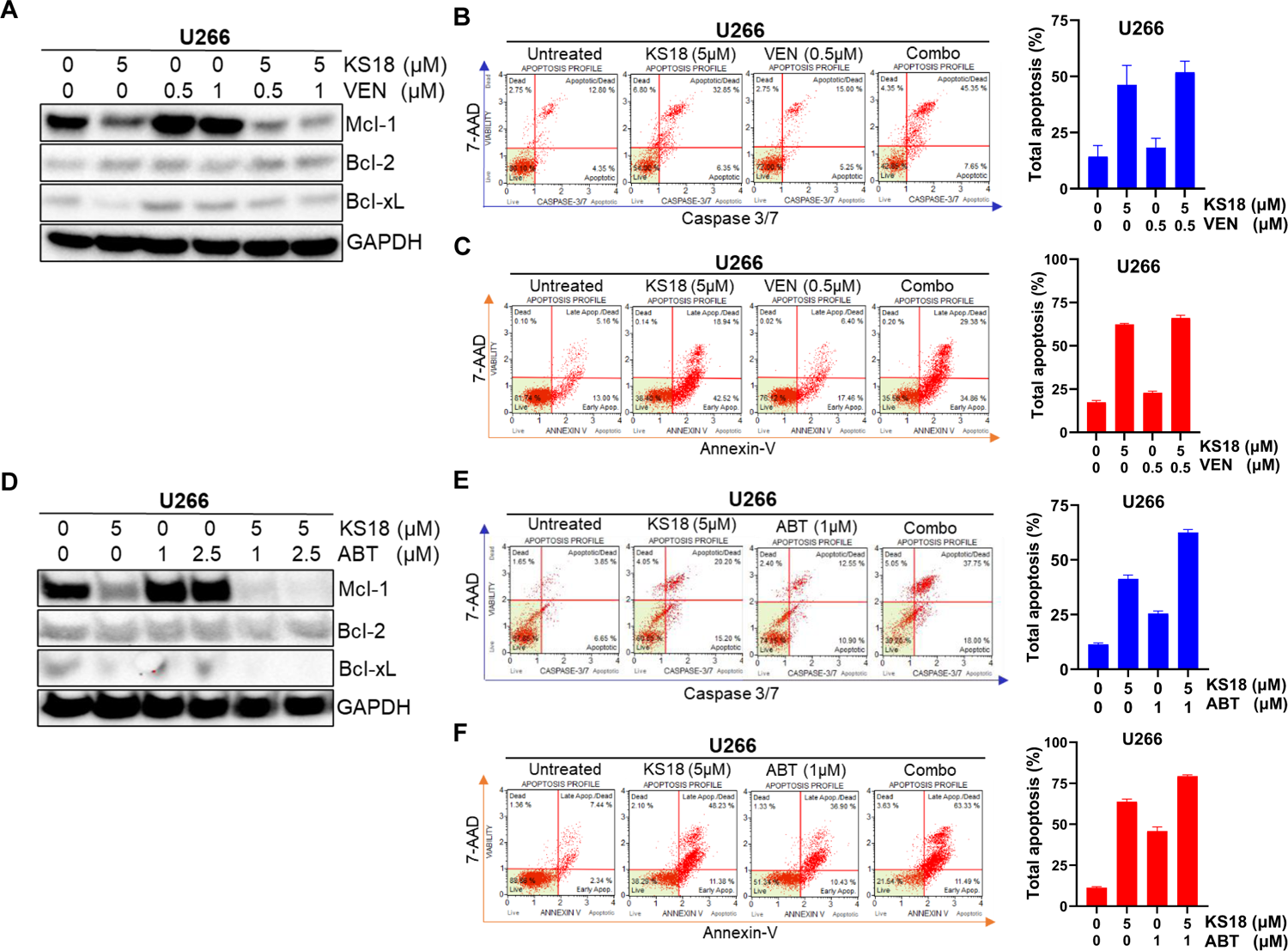
KS18 increases the efficacies of venetoclax and ABT-737 in MM cells. ***A,*** The effect of venetoclax (VEN) and KS18 on anti-apoptotic proteins. U266 cells were treated for 24h with VEN (0.5 and 1µM) alone and in conjunction with KS18 (5µM), and immunoblotting was performed against the specified antibodies. ***B&C,*** Caspases 3/7 and Annexin V staining are used to detect apoptotic cells. U266 cells were treated for 24h with VEN (0.5µM) alone or in conjunction with KS18 (5µM), then stained with caspases 3/7 dye or Annexin V dye and evaluated using the Muse® Cell Analyzer. Graphical representations were created using GraphPad prism. ***D,*** The effect of ABT-737 (ABT) and KS18 on anti-apoptotic proteins. U266 cells were treated for 24h with ABT (1 and 2.5µM) alone and in conjunction with KS18 (5µM), and immunoblotting was performed against the specified antibodies. ***E&F,*** Caspases 3/7 and Annexin V staining are used to detect apoptotic cells. U266 cells were treated for 24h with ABT (1µM) alone or in conjunction with KS18 (5µM), then stained with caspases 3/7 dye or Annexin V dye and evaluated using the Muse® Cell Analyzer. Graphical representations were created using GraphPad prism.

We investigated whether ABT-737 had similar effect as was observed with venetoclax. As expected, ABT-737 alone increased Mcl-1 expression, while KS18 and ABT-737 suppressed Mcl-1 (Fig. 4D; Supplementary Fig. 3E). The apoptosis-inducing ability of KS18 in conjunction with ABT-737 was confirmed (Fig. 4E-F; Supplementary Fig. 3F-G). As observed with venetoclax, the addition of KS18 (5µM) to ABT-737 reduced cell viability significantly in MM cells (Supplementary Fig. 3H). Our findings indicate that Mcl-1 inhibition is required to improve the therapeutic outcomes of venetoclax and ABT-737.

### KS18 re-sensitizes bortezomib-resistant MM cells to the drug and kills resistant MM cells

Because acquired resistance to standard care therapy is the leading cause of recurrence in MM, we investigated how effective KS18 is in MM-resistant cells. We created resistant cells to a variety of MM chemotherapeutic drugs (bortezomib, lenalidomide, venetoclax, and ABT-737) and discovered that all resistant cell lines expressed pro-survival Bcl-2 family proteins (Bcl-2, Bcl-xL, and Mcl-1) differently (Supplementary Fig. 4A-B). Nonetheless, the high expression of Mcl-1 compared to Bcl-2 or Bcl-xL suggests that Mcl-1 is a driver of resistance in these models. We investigated KS18’s efficiency in resistant cells and discovered that it was significantly more efficient in resistant cells than in parental lines (Fig. 5A). We then examined whether KS18 can inhibit Mcl-1 expression in resistant cells, and as shown in Fig. 5B, KS18 can decrease Mcl-1 expression. Furthermore, we investigated whether KS18 therapy may influence the dynamics of Mcl-1 and pro-apoptotic proteins in resistant cells. KS18 therapy modified Mcl-1: Bim and Mcl-1: Bax interactions in resistant cells, and restored Bax apoptotic role (Fig. 5C-D). Furthermore, we looked at KS18 activity in U266-bortezomib resistance (U266-BTZ-R) cell’s colony formation. In U266-BTZ-R cells, KS18 therapy reduced colony formation. Compared to untreated controls, the colonies were reduced by 92% (Fig. 5E). We then investigated whether combining KS18 with bortezomib is a viable option for treating bortezomib-resistant cells. The U266-BTZ-R cells were treated either alone or in combination with KS18 (5µM) or bortezomib (20nM). The addition of KS18 to the treatment combination increased the response to bortezomib in resistant cells, indicating that KS18-based treatment combinations are effective against resistant cells and can re-sensitize MM cells to chemotherapy (Fig. 5F). Our findings suggest a compelling rationale for employing KS18 to treat MM resistant cells.

**Figure 5:**
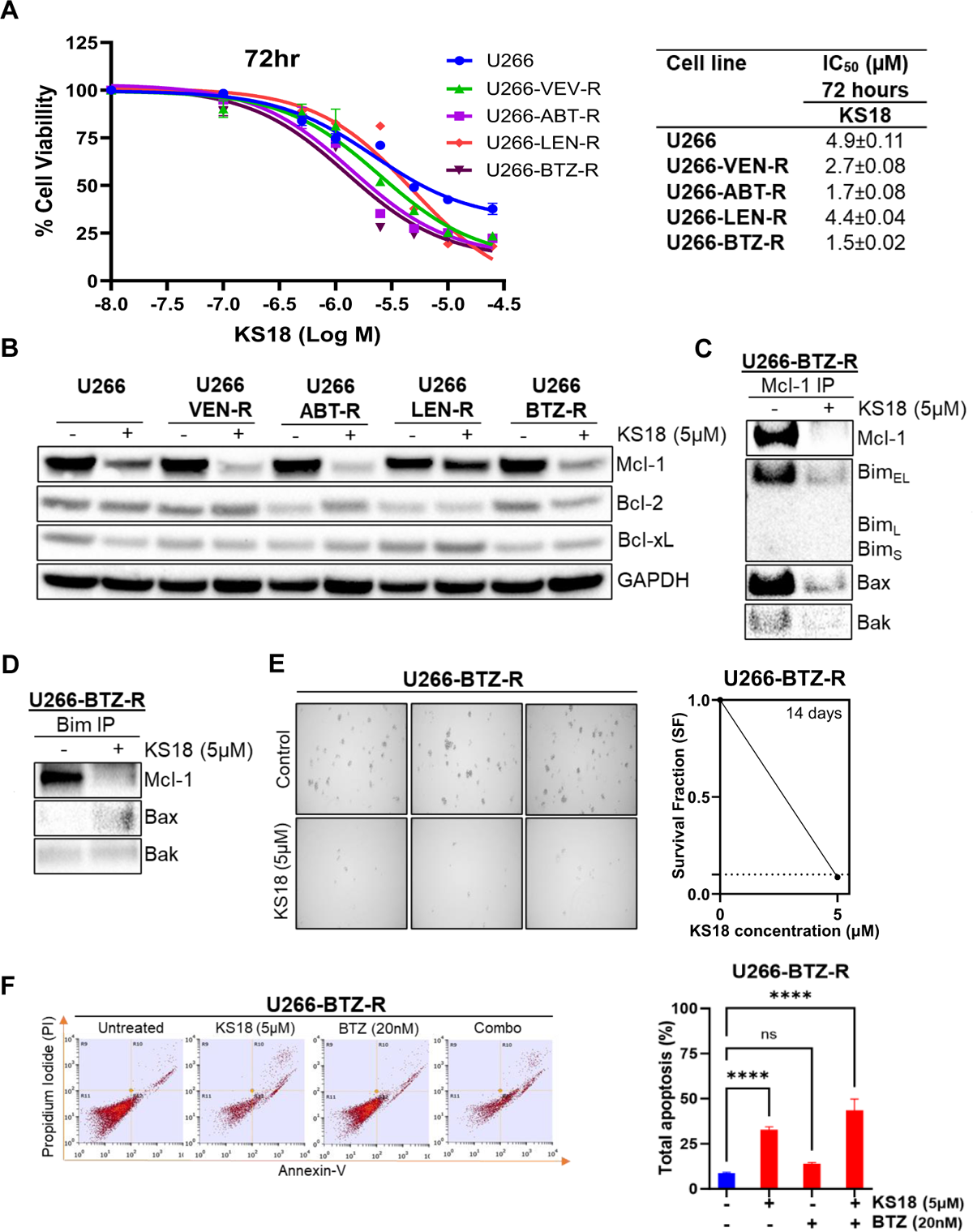
KS18 is cytotoxic against MM-resistant cells and re-sensitize MM-bortezomib resistant cells to bortezomib. ***A,*** A panel of human MM U266-resistant cell lines (U266-VTX-R, U266-ABT-R, U266-LEN-R, and U266-BTZ-R) were treated for 72h with increasing dosages of KS18 (0-25µM), and cytotoxicity was assessed using the MTT assay. GraphPad prism was used for graphical representation and IC_50_ calculation. ***B,*** KS18’s impact on anti-apoptotic proteins in susceptible and resistant cell lines. KS18 (5µM) was applied to several resistant cells for 24h, and the expression of anti-apoptotic proteins was assessed by immunoblotting. Anti-GAPDH was employed to quantify protein loading. ***C&D,*** U266 bortezomib resistant cells were treated with KS18 (5µM) for 24h before being lyzed. As specified in ‘MATERIALS AND METHODS’ section, the whole cell extract was immunoprecipitated with either Mcl-1 or Bim, and immunoblotting was done with the aforementioned antibodies. ***E,*** U266 bortezomib resistant (U266-BTZ-R) cells were grown in colony supporting matrix for 14 days, with and without KS18 at a concentration of 5µM. Following the visual counting of colonies with a microscope, the survival fraction (SF) was calculated in relation to the untreated control group, and a graphical representation was made using GraphPad prism. ***F,*** KS18 re-sensitizes U266-BTZ-R cells to bortezomib therapy. Annexin V labeling in flow cytometry was used to detect apoptotic cells. U266-BTZ-R cells were treated for 24h with BTZ (20nM) or KS18 (5µM) alone or in combination, then stained with Annexin V dye and evaluated by flow cytometry, and a graphical representation was created using GraphPad prism. **** P ≤ 0.0001; ns: non-significant.

### With its ability to enhance the venetoclax apoptotic response, KS18 shows promise as an adjuvant treatment for bortezomib-resistant MM cells

We wanted to explore the combination of KS18 and venetoclax in bortezomib resistant cells because Mcl-1 overexpression has been linked to resistance.^46,47^ We used various concentrations of venetoclax alone or in combination with KS18 (2.5µM) to treat U266-BTZ-R cells. As expected, venetoclax increased Mcl-1 levels; however, when combined with KS18, Mcl-1 levels were lowered, as were Bcl-2 and Bcl-xL (Fig. 6A). The combination of ABT-737 and KS18 had similar effects in U266-BTZ-R (Fig. 6B). According to a recent study, dexamethasone enhances the expression of both Bcl-2 and Bim in MM, which shifts Bim binding towards Bcl-2 and promotes Bcl-2 dependency in MM.^48^ Thus, dexamethasone makes MM-bortezomib resistant cells susceptible to venetoclax. We evaluated whether adding KS18 to a dexamethasone and venetoclax regimen would increase apoptosis; indeed, adding KS18 led in apoptosis in over 95% of MM-bortezomib resistant cells (Fig. 6C). A three-drug regimen of bortezomib, cyclophosphamide, and dexamethasone (VCd or CyBorD) is an essential therapy option for patients with recurrent MM who are unresponsive to lenalidomide and daratumumab. Furthermore, VCd is an appropriate alternative for patients who are at higher risk of lenalidomide problems (i.e., acute kidney failure, increased thromboembolic risk) and for countries where lenalidomide is not allowed for initial therapy.^41^ In this study, we put KS18 through its paces against bortezomib-resistant cells. Our findings verified the synergistic effect of co-treatment. In the KS18/cyclophosphamide/dexamethasone regimen, *in vitro* synergism was found (Fig. 6D). Vincristine, doxorubicin, and dexamethasone (Vad) is another combination used as induction therapy for newly diagnosed MM.^42^ We investigated the combination of KS18, doxorubicin, and dexamethasone. The KS18/doxorubicin/dexamethasone regimen demonstrated *in vitro* synergism (Fig. 6E). In bortezomib-resistant cells, we compared the efficacy of KS18 to that of other commercially available Mcl-1 inhibitors such as S63845, VU661013, and AZD5991. Mcl-1 inhibitors at various concentrations (0-2500 nM) were applied to bortezomib resistant cells for 72h. Surprisingly, KS18 outperformed other Mcl-1 inhibitors and demonstrated a better cytotoxic effect in bortezomib resistant cells (Supplementary Fig. 4C), indicating that while commercially available Mcl-1 inhibitors lose efficacy in resistant cells, KS18 remains effective in both sensitive and resistant cells. The effectiveness of Bcl-2/Bcl-xL inhibitors and KS18 against bortezomib-resistant cells was investigated in detail, and the results show that KS18 remains the most effective against these cells (Supplementary Fig. 4D). This suggests that among the resistant population, Mcl-1 inhibitors are a better option than Bcl-2/Bcl-xL inhibitors, particularly in Mcl-1 overexpressing phenotypes.

**Figure 6:**
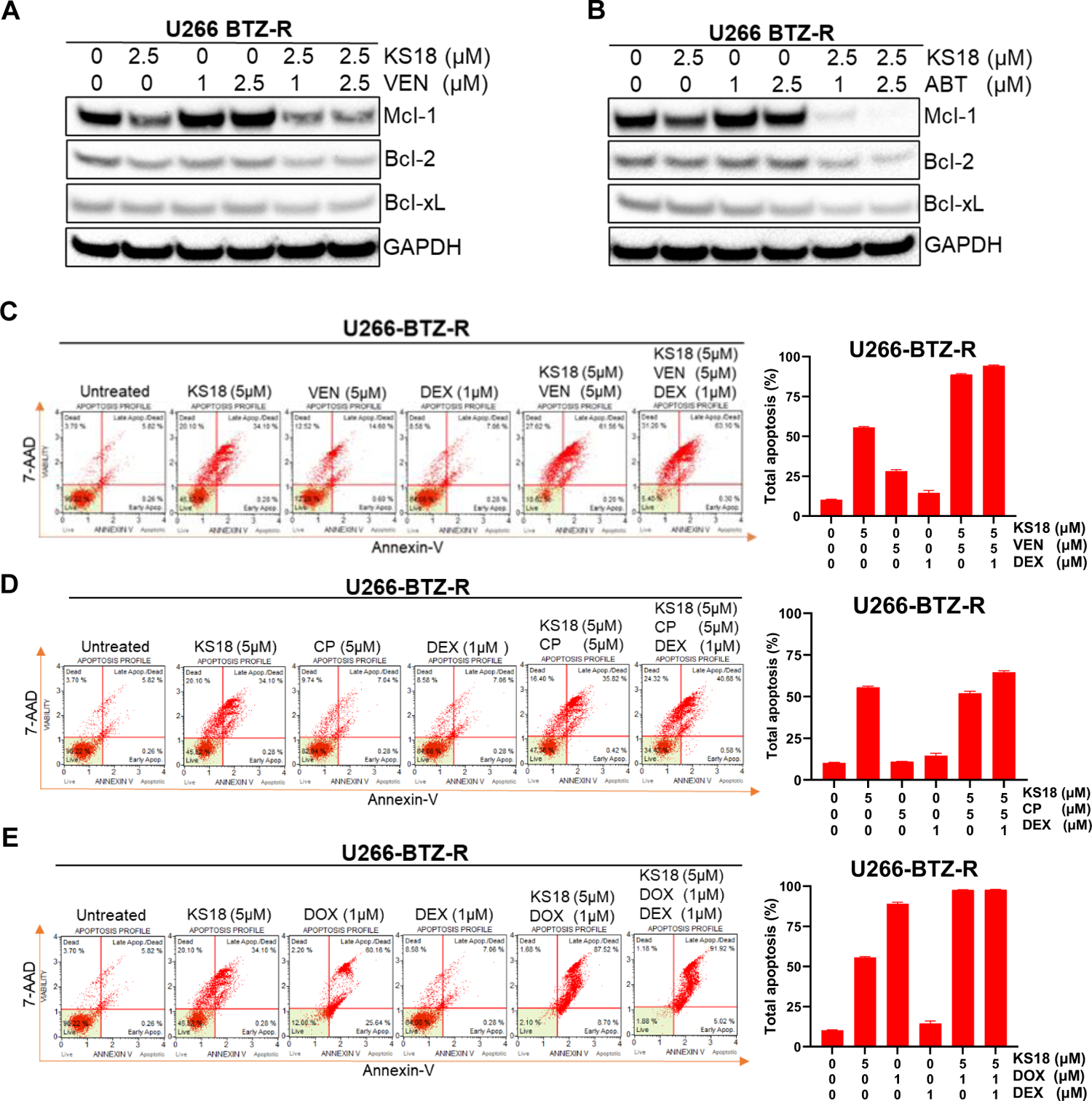
KS18 increases the efficacies of Bcl-2/Bcl-xL inhibitors in bortezomib resistant cells. ***A,*** U266-BTZ-R cells were treated with venetoclax (VEN) (1 and 2.5μM) alone and in combination with KS18 (5μM) for 24h, and immunoblotting was performed on anti-apoptotic proteins. ***B,*** U266-BTZ-R cells were treated with ABT-737 (ABT) (1 and 2.5μM) alone and in combination with KS18 (5μM) for 24h, and immunoblotting was performed on anti-apoptotic proteins. ***C,*** Annexin V staining to detect apoptotic cells. U266-BTZ-R cells were treated for 24h with KS18 (5μM) alone and in combination with VEN (5µM) and dexamethasone (DEX) (1µM), then stained with Annexin V dye and analyzed by the Muse Cell Analyzer and a graphical representation was created using GraphPad prism***. D,*** U266-BTZ-R cells were treated for 24h with KS18 (5μM) alone and in combination with cyclophosphamide (CP) (5µM) and dexamethasone (DEX) (1µM), then stained with Annexin V dye and analyzed by the Muse Cell Analyzer, and a graphical representation was created using GraphPad prism ***E,*** U266-BTZ-R cells were treated for 24h with KS18 (5μM) alone and in combination with doxorubicin (DOX) (1µM) and dexamethasone (DEX) (1µM), then stained with Annexin V dye and analyzed by the Muse Cell Analyzer, and a graphical representation was created using GraphPad prism.

We explored KS18 as an adjuvant therapy in venetoclax and ABT-737 resistant cells. We found that treating U266-venetoclax resistant cells with venetoclax and KS18 alone or in combination decreased the expression of Bcl-2, Bcl-xL, and Mcl-1 (Supplementary Fig. 4E). Similarly, we extended our observations with U266-ABT-737 resistant cells. Bcl-2, Bcl-xL, and Mcl-1 expression were also lowered by the combination of ABT-737 and KS18 (Supplementary Fig. 4F). Overall, these findings show that KS18 is an excellent Mcl-1 inhibitor and can be utilized as an adjuvant therapy in combination with bortezomib, dexamethasone, cyclophosphamide, doxorubicin, venetoclax, and ABT-737 in a range of resistant cells.

### KS18 shows promise in a model of MM xenografts as an anti-neoplastic agent

The MM xenograft model was developed by subcutaneous (S.C.) injection of U266 cells into the right flank of the NSG mice. The animals were randomized after approximately 10 days of tumor cell injection into three groups (when the tumor reached approximately 100mm^3^) (Fig. 7A). We tested KS18’s efficacy and safety in pilot tests and found that one weekly dose (q.wk) of 5 and 10 mg/kg was well tolerated (Fig. 7B). One mouse administered with 10 mg/kg was sluggish and hunched back at day 23, was euthanized. Importantly, all mice maintained normal weight compared to vehicle-treated mice (Fig. 7B). Similarly, we examined KS18’s effectiveness in MM xenografted mice. The NSG mice were treated with 5 and 10 mg/kg, q.wk. We found that two doses of KS18 considerably reduced tumor burden in mice (Fig. 7C). These data strongly imply we created a safe and a highly effective Mcl-1 inhibitor.

**Figure 7:**
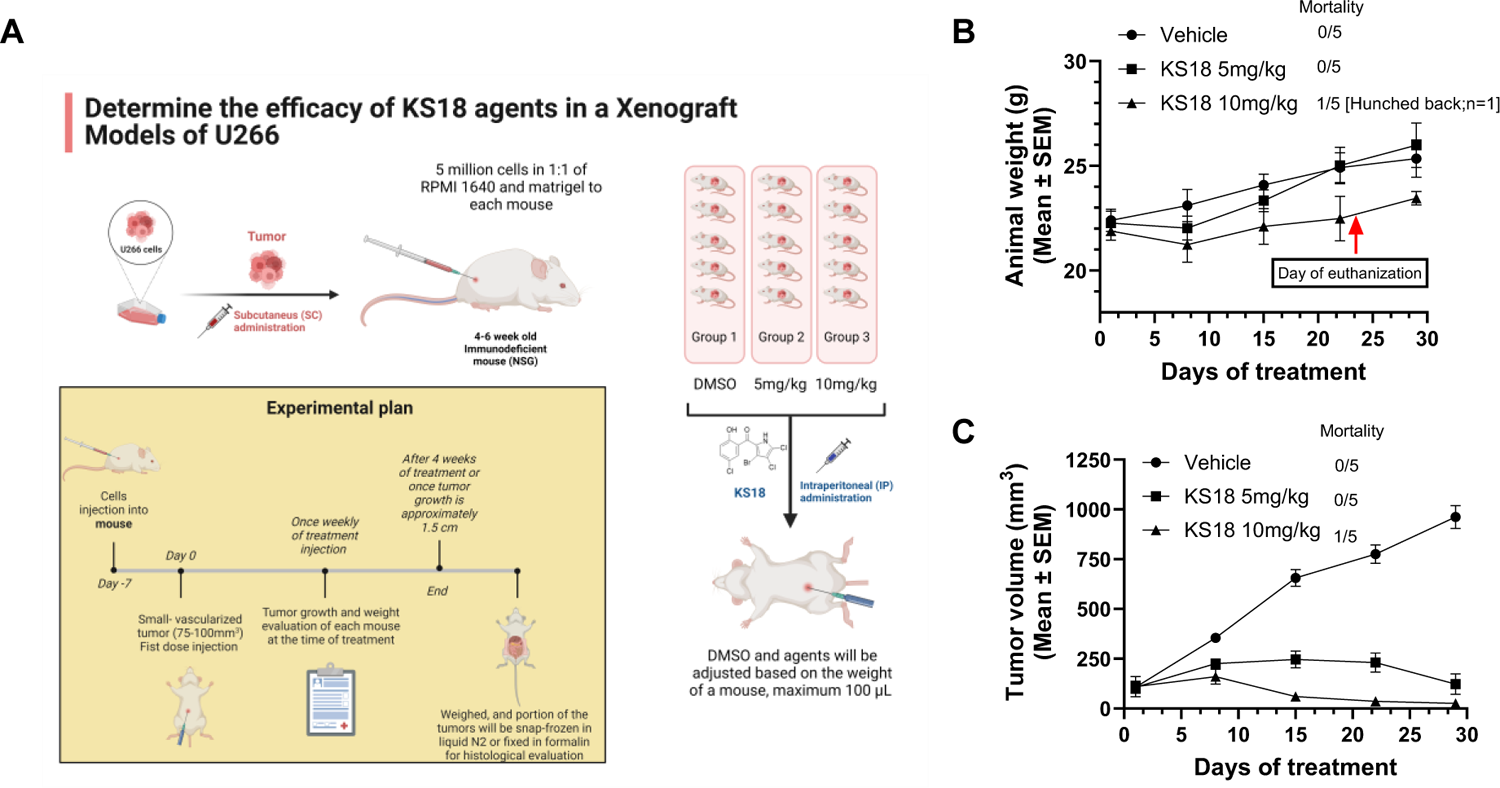
KS18 is safe and inhibits MM tumor growth. ***A,*** in a 1:1 mixture of RPMI 1640 medium and matrigel, five million cells were injected subcutaneously (SC) in the right flank of mice. Once tumor size reached approximately 100mm^3^, mice were randomly separated into three groups and treated with either DMSO or KS18 (5 and 10 mg/kg) intraperitoneally (IP) once a week (q.wk) for four weeks. ***B,*** KS18 is nontoxic to mice. Two KS18 doses were administered over the course of 28 days, and animal weights in xenografted mice were measured. On day 23, one sluggish mouse was euthanized. Presentation of animal body weight as mean ± SEM for vehicle (DMSO) and KS18 treated mice. ***C,*** Graph depicts tumor volume in mice treated with vehicle or KS18.

## DISCUSSION

MM is difficult to cure, and recent breakthroughs in treatment have not changed the reality that it is an incurable disease; all patients eventually relapse or become resistant. ^3,7^ Members of the Bcl-2 family play critical roles in the process of mediating the intrinsic apoptotic pathway. Importantly, overexpression of Mcl-1 gene is linked to a poor prognosis and resistance to treatment.^12,13^ Our findings show that KS18 specifically inhibits Mcl-1. KS18 stimulates Mcl-1 phosphorylation at Ser159/Thr163 and ubiquitination, resulting in a significant decrease in Mcl-1 protein levels, showing that KS18 activates the ubiquitin/proteasome-dependent protein degradation system (UPS). Importantly, KS18 treatment has a remarkable potential to induce complete apoptosis in cells and MM patient samples in a caspase-dependent manner. Our findings further show that KS18 activates caspase-3 in MM cells by activating the MOMP pathway, which includes Bak and Bax and leads in the cytoplasmic release of cytochrome C. KS18 also works in tandem with bortezomib and venetoclax to increase apoptosis in MM cells.

The cytotoxic potential of KS18 was evaluated and compared to that of currently available chemotherapeutic medications. According to our findings, KS18 significantly reduced the viability of MM cell lines and was more effective than most currently used chemotherapies. Furthermore, we discovered that adding a low dose of dexamethasone to the KS18/bortezomib regimen magnified the efficacy of the bortezomib/dexamethasone (Vd) regimen and forms an excellent alternative to the bortezomib/lenalidomide/dexamethasone (VRd) regimen, making it an intriguing contender for further investigation as a potential therapeutic option for MM.

One of the study’s key findings is that Mcl-1 overexpression contributes to developing treatment resistance in MM cells. This is supported by the fact that all MM-resistant cell lines tested in this study, including those resistant to bortezomib and venetoclax, have Mcl-1 overexpression. Given the increasing reliance of resistant MM cells on Mcl-1 overexpression, we reasoned that targeting Mcl-1 was a reasonable strategy for treating resistant MM and could effectively overcome drug resistance. In MM-bortezomib resistant cells, KS18 outperforms venetoclax and commercially available Mcl-1 inhibitors such as S63845, VU661013, and AZD5991, with an average IC_50_ value of 1.5µM. KS18 not only restores apoptosis in bortezomib-resistant MM cells, but it also works in conjunction with venetoclax to increase apoptosis. This is an important finding. KS18 has been proven *in vitro* to overcome bortezomib resistance and re-sensitize MM cells to the effects of bortezomib, marking a potential step forward in treating the disease. Compared to untreated controls, KS18 treatment reduced colony formation in bortezomib-resistant MM cells by 92%. *In vivo* investigations on MM mouse models add to the evidence that KS18 is both effective and safe. Mice showed significant tumor shrinkage after four weeks of therapy with a single acceptable dose per week, indicating this compound’s powerful anti-myeloma effects.

Finally, the study underlines the role of Mcl-1 in MM and drug resistance development, as well as the potential utility of KS18 as a treatment for newly diagnosed MM patients and those who have developed resistance to chemotherapy. Our results reveal that KS18 is a highly specific and potent inhibitor of Mcl-1. KS18 is more effective than other chemotherapy agents and increases the potency of other chemotherapeutic treatments, such as bortezomib and venetoclax, against MM cells. Furthermore, our findings show that KS18 may be effective as an adjuvant to therapies aimed at overcoming bortezomib resistance, re-sensitizing MM resistant cells to chemotherapy, and amplifying the response to venetoclax treatment. More research with animal models and sensitive- and resistant-MM patient samples with varied genetic lesions is required to direct the remarkable potential of this innovative medicine to human treatment trials to assess the safety and efficacy of KS18.

## ACKNOWLEDGEMENT

The Camden Research Initiative Fund (M.K.P, T.B.A, and S.C.J) and an interdepartmental fund from Cooper Medical School of Rowan University, Camden, NJ (M.K.P) sponsored this work. The synthesis of the analogs was partially funded by National Institutes of Health Grant ES028244, which was sub-awarded to (S.G.A and K.G).

## AUTHORSHIP

Contribution: O.S.O designed, performed experiments, analyzed the data, and wrote the manuscript; S.G.A and K.G. conceived ideas, contributed to providing essential reagents and synthesis of the KS18, and critically evaluating the study; S.K.S. designed and preformed docking studies and critically revised the manuscript. M.K.P., T.B.A., and S.C.J. conceived ideas, designed the study, interpreted the results, and revised the manuscript. Conflict-of-interest disclosure: The authors declare no competing financial interests.

## SUPPLEMENTARY FIGURE LEGENDS

**Supplementary Figure 1:**
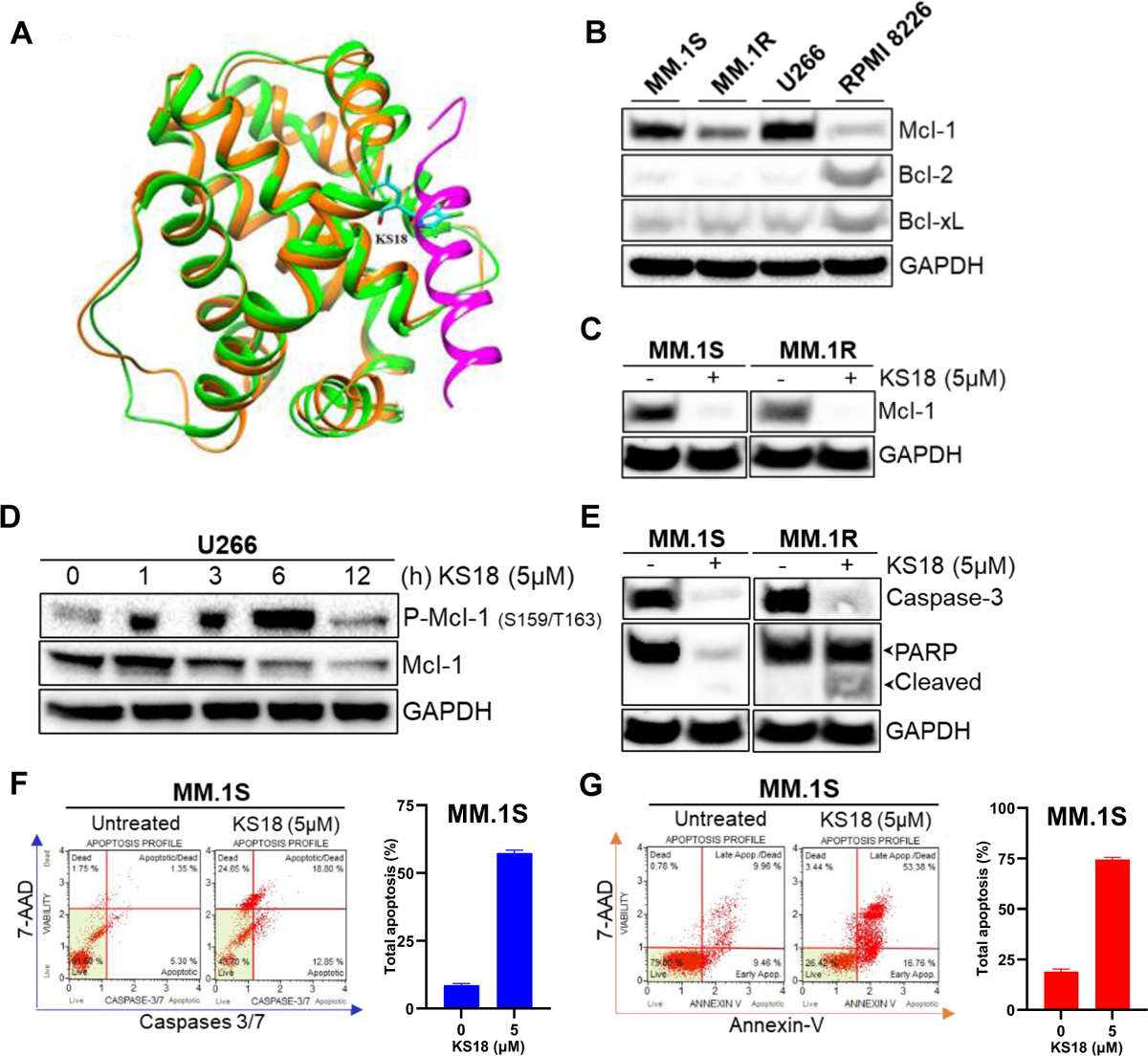
KS18 inhibits Mcl-1 and induces apoptosis in MM cells. ***A.*** Superimposition of Mcl-1 (orange)-Noxa (magenta) in complex with KS18 (stick cyan) with Mcl-1. ***B.*** Myeloma cells show differential expression of anti-apoptotic Bcl-2 family protein. ***C,*** The effect of KS18 on Mcl-1 in two MM cell lines. ***D,*** Time-dependent effect (0-12h) of KS18 (5μM) on the phosphorylation of Mcl-1 in MM cells. ***E,*** The effect of KS18 (5μM) on caspase-3 and PARP proteins in various MM cells. For sections ***B-E***, MM cells treated with or without KS18 (5µM) for either specified time (section D) or 24h. Following incubation, cells were collected, lyzed, and immunoblotting for the aforementioned antibodies was done. The anti-GAPDH blotting was performed to determine loading. ***F&G,*** Caspases 3/7 and Annexin V staining were used to identify apoptotic cells. MM.1S cells were treated with KS18 (5µM) for 24h before being stained with caspases 3/7 dye or Annexin V dye as described in ‘MATERIALS AND METHODS’ and evaluated using the Muse® Cell Analyzer. The total number of apoptotic cells was enumerated, and a graph was created using the GraphPad prism software.

**Supplementary Figure 2:**
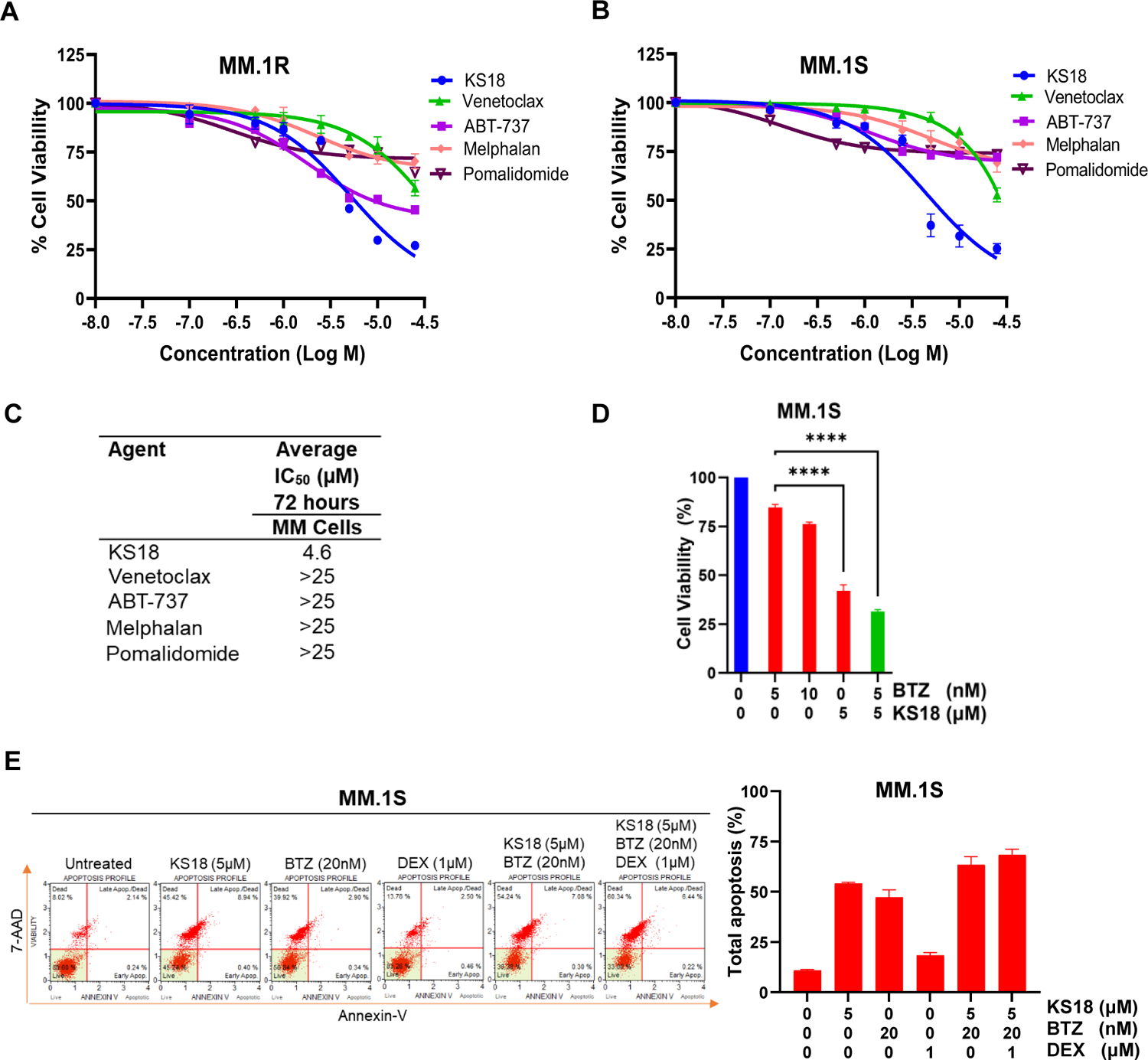
KS18 is more potent than other chemotherapeutic agents and increases the efficacy of bortezomib in MM cells. ***A-C,*** Cell viability was evaluated using the MTT test after 72h of treatment with increasing dosages (0-25µM) of KS18 and other chemotherapeutic drugs (venetoclax, ABT-737, melphalan, and pomalidomide). GraphPad prism was used for graphical representation and IC_50_ calculation. ***D***, Bortezomib (BTZ) (5 and 10nM) alone and BTZ (5nM) in combination with KS18 (5µM) were used to treat MM.1S cells, and cell viability was determined using the MTT test. GraphPad prism was used to construct graphical representations and statistical analyses. ***E***, Apoptotic cells are detected by Annexin V staining. MM.1S cells were treated for 24h with KS18 (5µM) alone or in combination with BTZ (20nM) and DEX (1µM), then stained with Annexin V dye and evaluated using the Muse® Cell Analyzer. The total number of apoptotic cells was counted and graphed using the GraphPad prism software.

**Supplementary Figure 3:**
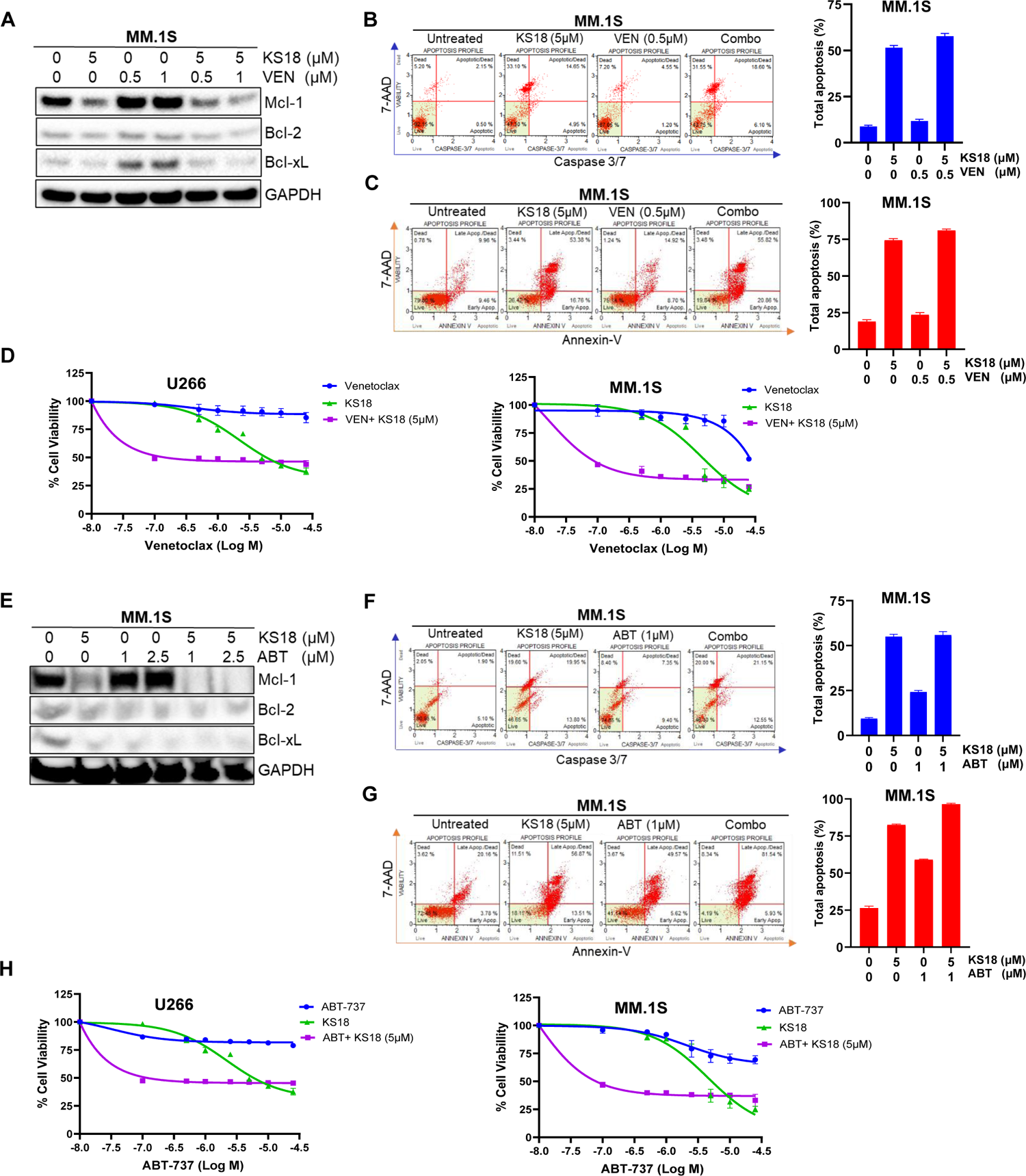
KS18 increases the efficacies of venetoclax and ABT-737 in MM cells. ***A,*** Venetoclax (VEN) therapy increases Mcl-1 protein expression in MM.1S cells while KS18 suppresses it. For 24h, MM.1S cells were treated with VEN (0.5 and 1µM) alone or in combination with KS18 (5µM), and immunoblotting was conducted. ***B&C,*** Caspases 3/7 and Annexin V labeling were used to detect apoptotic cells. MM.1S cells were treated for 24h with VEN (0.5µM) alone or in conjunction with KS18 (5µM), then labeled with caspases 3/7 or Annexin V dye and evaluated using the Muse Cell Analyzer. GraphPad prism software was used to count and graph the apoptotic cells. ***D,*** U266 and MM.1S cells were treated for 72h with increasing dosages (0-25µM) of VEN alone or in combination with KS18 (5µM), and cell viability was assessed using the MTT test and GraphPad prism. ***E,*** Treatment with ABT-737 (ABT) produces Mcl-1 protein in MM.1S cells, while KS18 greatly inhibits this induction and increases ABT efficacy. For 24h, MM.1S cells were treated with ABT (1 and 2.5µM) alone or in conjunction with KS18 (5µM), and immunoblotting was conducted. ***F&G,*** Caspases 3/7 and Annexin V labeling were used to detect apoptotic cells. MM.1S cells were treated for 24h with ABT (1µM) alone or in conjunction with KS18 (5µM), then stained with caspases 3/7 dye or Annexin V dye and evaluated using the Muse® Cell Analyzer. The total number of apoptotic cells was counted and graphed using the GraphPad prism software. ***H,*** U266 and MM.1S cells were treated for 72h with increasing dosages (0-25 µM) of ABT alone or in combination with KS18 (5 µM), and cell viability was evaluated using the MTT assay and graphed using the GraphPad prism program.

**Supplementary Figure 4:**
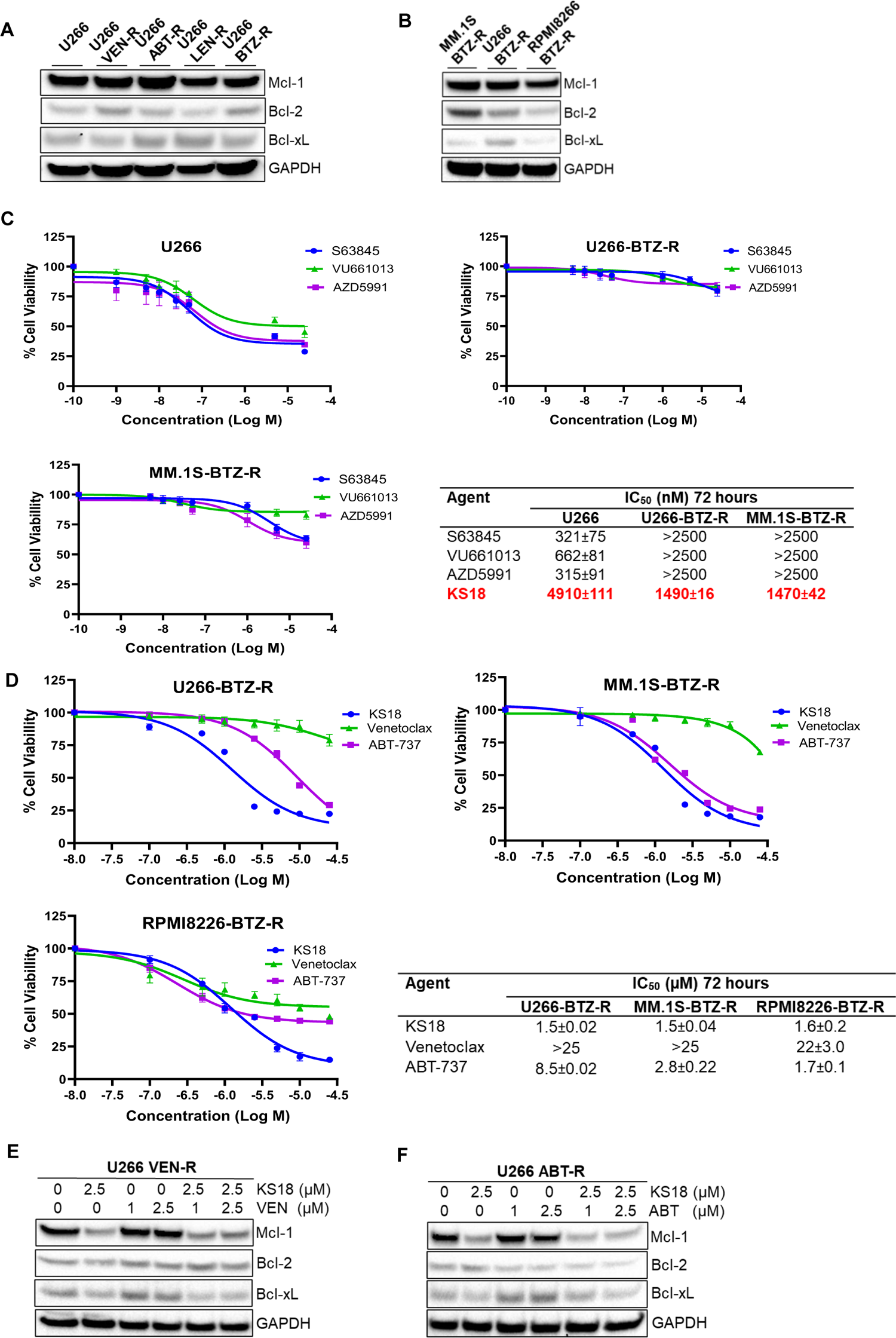
KS18 outperformed other Mcl-1 inhibitors in MM-bortezomib resistant cells. ***A&B,*** Mcl-1 and other anti-apoptotic proteins expression in MM-resistant cell lines determined by western blot. ***C,*** Cell survival was evaluated using the MTT test after 72h of treatment with increasing dosages (0-2500nM) of three selective Mcl-1 inhibitors (S63845, VU661013, and AZD5991). GraphPad prism was used to construct the graphical depiction and the IC_50_ calculation. ***D,*** A panel of human MM bortezomib-resistant cell lines (MM.1S-BTZ-R, U266-BTZ-R, and RPMI8226-BTZ-R) were treated for 72h with increasing dosages (0-25µM) of KS18, venetoclax (VEN), and ABT-737 (ABT), and cell viability was determined using the MTT assay. GraphPad prism was used to construct the graphical depiction and the IC_50_ calculation. ***E,*** U266-VTX-R cells were treated with VEN (1 and 2.5μM) alone and in combination with KS18 (2.5μM) for 24h, and immunoblotting was performed. ***F,*** U266-ABT-R cells were treated with ABT (1 and 2.5μM) alone and in combination with KS18 (2.5μM) for 24h, and immunoblotting was performed.

## Notes

### Competing Interest Statement

The authors have declared no competing interest.

